# Improved structural modelling of antibodies and their complexes with clustered diffusion ensembles

**DOI:** 10.1101/2025.02.24.639865

**Authors:** Marco Giulini, Xiaotong Xu, Alexandre MJJ Bonvin

## Abstract

**Motivation:** Gaining structural insights into antibody-antigen complexes is crucial for understanding antigen recognition mechanisms and advancing therapeutic antibody design. However, accurate prediction of the structure of highly variable complementarity-determining region 3 on the antibody heavy chain (CDR-H3 loop) remains a significant challenge due to its increased length and conformational variability. While AlphaFold2-multimer (AF2) has made substantial progress in protein structure prediction, its application on antibodies and antibody-antigen complexes is limited by the weak evolutionary signals in the CDR region and the lack of structural diversity in its output.

**Results:** To address these limitations, we propose a workflow that combines AlphaFlow to generate ensembles of potential loop conformations with integrative modelling of antibody-antigen complexes with HADDOCK. Improving the structural diversity of the H3 loop increases the success rate of subsequent docking tasks. Our analysis shows that while AF2 generally predicts accurate antibody structures, it struggles with the H3 loop. In cases where AF2 mispredicts the loop, we leverage AlphaFlow to generate ensembles of loop conformations via diffusion-based sampling, followed by clustering to produce a structurally diverse set of models. We demonstrate that these ensembles significantly improve antibody-antigen docking performance compared to the standard AF2 ensembles.

**Availability and implementation:** The input datasets and codes involved in this research are available at https://github.com/haddocking/alphaflow-antibodies. All the resulting modelling data are available from Zenodo (https://zenodo.org/records/14906314).

## 1 Introduction

Antibodies are large, Y-shaped proteins consisting of two light and two heavy chains, produced by the immune system that bind to pathogenic molecules (antigens) with high specificity. The high specificity and designability of antibodies have made them ideal molecules for therapeutic purposes, with more than 170 antibodies being approved by the US FDA as of 2023.

Gaining structural insights into antibody-antigen complexes is vital for understanding their functions and optimizing antibody design. To compensate for the high costs of wet-lab experiments, in-silico approaches are increasingly used to model and predict how antibodies interact with their specific antigens. Antibodies bind their cognate antigen by means of their Complementarity Determining Regions (CDRs), namely six highly variable loops, three from the light and three from the heavy chain, located at the end of the two arms of the Y-shaped structure. Predicting the conformation of such loops is challenging due to the sheer size of their sequence space and the lack of coevolution information. Due to its unique position in the genome at the intersection of three different gene loci, the third loop on the heavy chain (H3) stands out as the most challenging, being much longer and more heterogeneous than the other five loops, which show a limited number of different conformational clusters (North *et al*., 2011; Weitzner *et al*., 2015). It is estimated that 75% of H3 loops do not have a sub-Angstrom structural neighbour in the non-antibody world (Regep *et al*., 2017).

The recent abundance of available antibody structures in the Protein Data Bank (Berman *et al*., 2000) (9080 as of 31-12-2024, with more than 50% deposited in the last five years (Dunbar *et al*., 2014)), together with the development of general (Baek *et al*., 2021; Jumper *et al*., 2021) and antibody-specific (Ruffolo *et al*., 2023; Abanades *et al*., 2023) machine learning-based structure prediction methods, boosted the accuracy of antibody structure prediction. Nonetheless, the prediction of the correct H3 conformation remains challenging, with a consistent fraction of loops being mispredicted (Wu *et al*., 2022).

In this work, we first evaluate the performance of AlphaFold2-multimer (AF2) on the antibody prediction task. It is well-established (McCoy *et al*., 2024) that AF2 is one of the most accurate predictors for this purpose, though it has the limitation of providing limited model diversity in its output. To address this issue, several methods have been proposed to increase AlphaFold2’s output diversity (Wallner, 2023; Raouraoua *et al*., 2024; Wayment-Steele *et al*., 2024; Jing *et al*., 2024; Stein and Mchaourab, 2022), most of which involve modifications to the input Multiple Sequence Alignments (MSAs). For example, Wayment-Steele *et al*. showed that clustering MSAs by sequence similarity enable AF2 to generate high confidence alternative structural states for a known ‘fold-switching’ protein. Additionally, MassiveFold (Raouraoua *et al*., 2024) enhances AF2’s model diversity by repeatedly running AF with varying input parameters and models. Recently, AlphaFlow (AFL) (Jing *et al*., 2024) has tackled this challenge from a new perspective by leveraging diffusion models. AFL transforms AF2 from a regression model into a sequence-conditioned generative model, utilizing a custom iterative denoising framework and a polymer-structured prior distribution, combined with a scale-invariant noising process. This approach enables AlphaFlow to directly sample an infinite number of structures during inference.

Here we hypothesize that AlphaFlow can be efficiently used to increase the structural diversity of the predicted H3 loops of antibodies, enabling the exploration of more conformations compared to the standard AlphaFold2-multimer pipeline, especially when the loop confidence is low (as measured by the predicted lDDT values).

By examining 54 antibodies released after the cutoff date of AF2’s training data, we show that, for difficult cases, predicting the structure of the antibody heavy chain with AFL increases the likelihood of obtaining a more accurate prediction with respect to AF2. Clustering these structures results in a highly heterogeneous antibody ensemble, which can be effectively used in downstream tasks such as antibody-antigen docking. We subsequently demonstrate how a docking protocol starting from the AFL ensemble outperforms the same protocol using the standard AF2 ensemble. We end by discussing the performance of AlphaFold3 (Abramson *et al*., 2024), which was released during the course of this project, in predicting the H3 loop conformation.

## 2 Methods

### 2.1 Dataset

For the dataset we consider the antibodies used in a previous study (Giulini *et al*., 2024), excluding those that were present in the training set of AlphaFold2-multimer version 2.3.1. This reduces the size of the dataset to 54 complexes (see SI Table1. for details).

### 2.2 Model details about AlphaFlow

AlphaFlow is a generative model built upon AlphaFold2, designed to produce conformational ensembles directly at inference time. This model uses custom flow matching and is tailored to fit the architecture and training objectives of AlphaFold. For our diffusion ensemble, we utilized the “PDB_base” version of AlphaFlow, which is trained on PDB structures to accurately model experimental ensembles derived from X-ray crystallography and cryo-EM under different conditions. Generating a single-chain diffusion model from an input sequence takes an average of around 40 seconds on a GeForce GTX 1080 GPU.

### 2.3 Construction of input ensembles for docking

We limited our dataset to the antibodies for which AlphaFold2’s H3 loop pLDDT is below 80 (see Results section 3.1 “AlphaFold antibody modelling” for details). For this “difficult” (LOW-80) dataset, we generate 1000 heavy chain antibodies using AlphaFlow in “PDB base” models. Additionally, we produce 25 predictions using our local installation of AlphaFold 2.3 with 5 random seeds, keeping all other parameters as default. To reduce the number of starting structures for subsequent docking, we cluster the 25 predictions in an ensemble of 5 models. An analysis of the quality of the H3 loop as a function of the number of models considered for clustering shows that 100 models are sufficient to keep a good quality of the H3 loop while reducing the computing time (see Results). These ensembles of 100 models are further clustered into 20 clusters, with the center of each cluster being considered for docking. This effectively reduces the number of AFL conformations by a factor of 5.

Next, we combine the generated heavy chains with predicted light chains from AF2 to create full antibody model ensembles. We observed several backbone clashes between the diffused H3 loop and the light chains during this process, as measured by an interatomic distance lower than 63% of the sum of the interatomic radii. Accordingly, we removed the structures showing more than one backbone clash. An energy minimisation step was conducted by running the “emref” module of HADDOCK3 on the remaining models. The energy-minimized antibody ensembles are used as starting models for docking.

### 2.4 HADDOCK Docking protocol

We use HADDOCK3 (Dominguez *et al*., 2003) (https://github.com/haddocking/haddock3, unpublished), our information-driven docking software, to perform the docking of the antibody ensembles to the desired antigen. Interface information is provided to HADDOCK3 using ambiguous interaction restraints, in which a set of input amino acids ideally should (active residues) or can (passive residues) be at the interface of the docked molecules. The workflow, defined in the HADDOCK3 configuration file, consists of six sequential steps: topology generation (topoaa), rigid-body docking (rigidbody), selection of the top ranked 200 models (seletop), semi-flexible refinement of the interface (flexref), energy minimisation (emref), and Fraction of Common Contacts clustering (clustfcc). This is equivalent to the standard HADDOCK2.4 (Honorato *et al*., 2024) pipeline. The corresponding input files and workflow configuration files are available from https://github.com/haddocking/alphaflow-antibodies. An example HADDOCK3 configuration file for docking is shown in the Supplementary Information.

## 3 Results

### 3.1 AlphaFold antibody modelling

First, we analysed the performance of AlphaFold2-multimer in modelling the 54 antibodies in our dataset (see Methods). Fig. 1a) shows the distribution of loop accuracies for the best ranked model of each structure (measured by the loop RMSD calculated upon aligning the structures over the framework region). This clearly highlights that the H3 loop is the only one for which the average prediction is more than 1.5Å away from the experimental loop conformation. In 31.5% of the dataset, the best-ranked AF2 model has a H3 loop more than 3 Å away from the reference structure.

**Figure 1.**
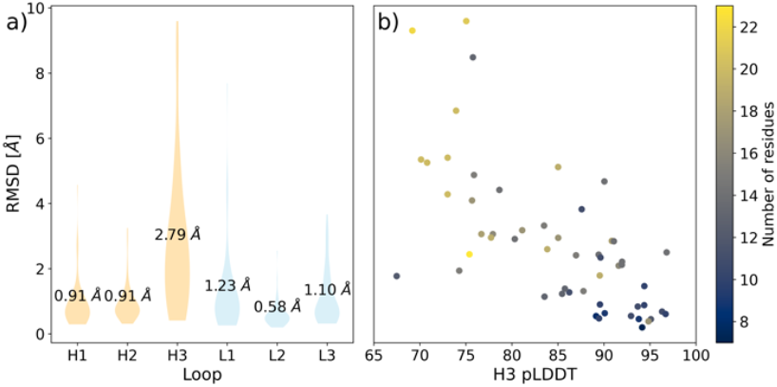
**a)** Violin plot of loop accuracy (measured by the loop RMSD from the reference crystal structure after superimposition on the framework region) over the six antibody hypervariable loops for the best ranked AlphaFold2 model. Mean values are reported for each loop. H3 stands out as the most difficult loop to be modelled by AF2. **b)** Scatter plot of the highest H3 pLDDT versus the corresponding H3-RMSD value. A clear anti-correlation exists between the two (Pearson R=-0.67), especially in the right region of the figure, where high values of H3-pLDDT almost always correspond to good accuracy. The scenario is different for lower pLDDTs, where the H3-RMSD can be either low or high at the same pLDDT value. The circles are colour-coded according to the length of the H3 loop.

The results are almost identical if, instead of considering the best-ranked AF2 model, we select the RMSD values of the loop with the highest H3 loop pLDDT (see SI Fig. 1).

Following insights from previous literature (Giulini *et al*., 2024; Kenlay *et al*., 2024), we checked how the H3 RMSD is related to the pLDDT calculated only over the H3 loop (Fig. 1b) considering the best ranked AF2 model in terms of H3 pLDDT. The two values are quite strongly anti-correlated, with a Pearson correlation coefficient equal to −0.67. This indicates that the H3 pLDDT is a strong predictor of loop accuracy. From this analysis, we assume that, if a H3 pLDDT higher than 80 is observed, the AF2 modelling is successful and we do not need to explore alternative strategies to sample its conformation.

For those antibodies whose best H3 loop has been predicted with pLDDT values lower than 80, some ambiguity arises, as the predicted structure can be quite accurate or very poor. In the remaining we focus on those 17 cases, which we name LOW-80. We further divide this dataset into two subsets, namely the 9 antibodies for which the best AF2 prediction is further than 3.0 (i.e., “Wrong”) from the reference structure (LOW-80-W) and 8 structures for which AF2 can generate accurate models (i.e., “Right”) (best H3 RMSD < 3.0Å) (LOW-80-R).

### 3.2 AlphaFlow performances for difficult loops

We run AlphaFlow (see Methods) on the heavy chain sequence (as AlphaFlow cannot handle multiple chains) of the 17 antibodies part of the LOW-80 dataset, generating ensembles of 1000 predictions per antibody. We calculated the models accuracy with respect to the reference heavy chain structure.

The most accurate models produced by AlphaFlow are on average 1.53 ±1.87Å better than the original ones generated by AF2 on this dataset. Fig. 2 reports the distribution of H3 RMSD. In the inset, a scatter plot of the AF2 versus AFL lowest H3 RMSD values is shown. AFL dramatically improves the quality of the loops for which AF2 was not accurate, with an average improvement of 2.74±1.75Å for the 9 antibodies with AF2 best H3 backbone RMSD > 3.0Å (LOW-80-W set). In contrast, AFL has hardly any impact over the other 8 cases (LOW-80-R set), for which the average improvement is negligible (0.16±0.70Å). AFL cannot improve structures in which the correct fold of the H3 loop has already been identified, but its usage is powerful where the conformation of the loop has been mispredicted. Understanding how to discriminate between these two cases with a predictor more accurate than pLDDT would be very useful.

**Figure 2.**
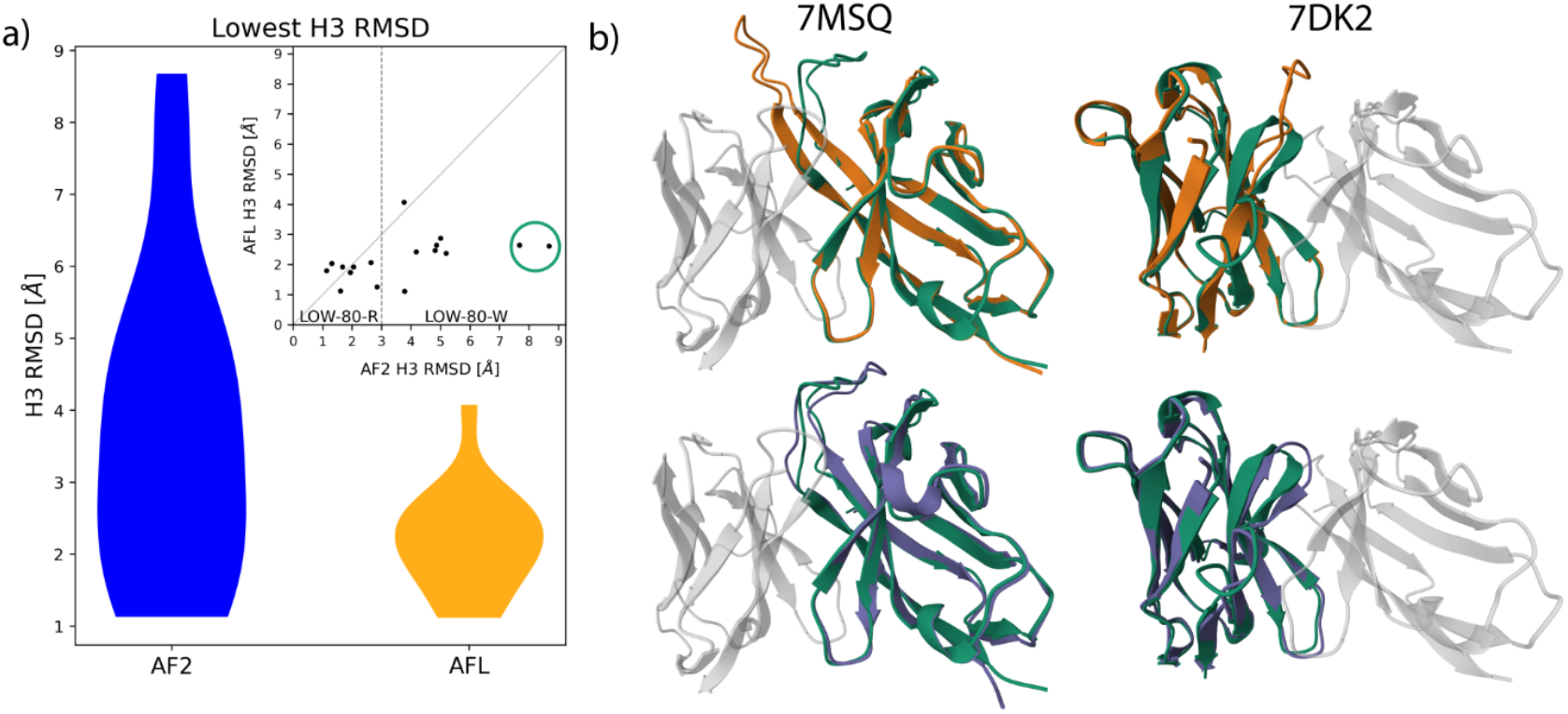
**a)** Violin plots of the distribution of lowest H3 RMSD values over the LOW-80 dataset for AlphaFold2 (25 generated models per antibody, left) and AlphaFlow ensembles (1000 generated models per antibody, right). AlphaFlow is always able to generate at least one model whose loop has a H3-RMSD < 3.0Å. **Inset:** Scatter plot of AFL vs AF2 H3 minimum RMSD values. Any point below the diagonal indicates an improvement in the H3 loop RMSD compared to the original AF2 model. The vertical line separates antibodies for which AF2 provides at least one model with H3-RMSD < 3.0A (LOW-80-R subset) from those structures for which all the AF2 models lie more than 3.0Å away from the ground truth (LOW-80-W subset). **b)** Superimposition between reference heavy chain structure (in green) and best Alphafold2 (orange) and best AlphaFlow (purple) models for two antibodies with exceptionally hard-to-predict H3 loops.

**Figure 3.**
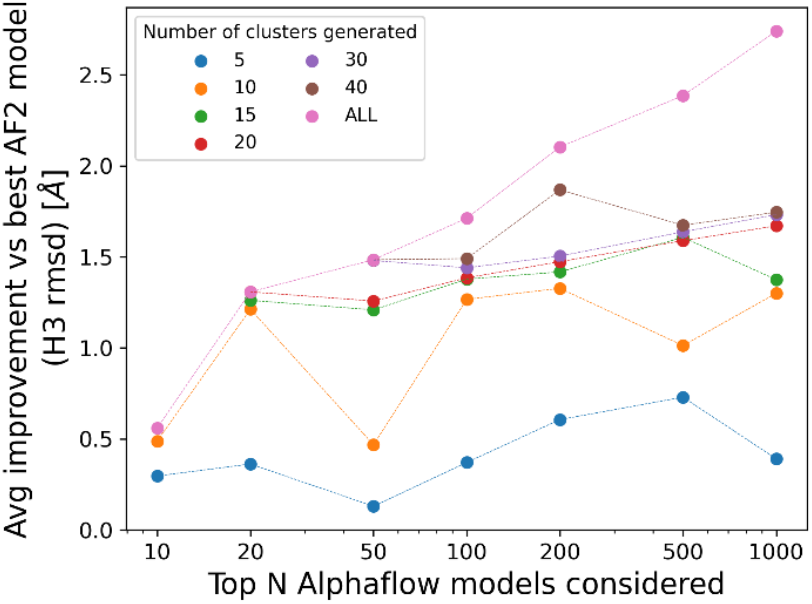
Clustering analysis of AlphaFlow heavy chain structures. The y-axis reports the H3 RMSD improvement observed with respect to the best structure of the AF2 ensemble, the x-axis corresponds to the number of AFL models that have been considered. Different colors correspond to different numbers of clusters. The ALL label denotes the best model across the considered AFL ensemble prior to clustering.

Generating 1000 models with AFL comes at non-negligible computational costs, roughly equal to generating 250 models with AF2. We therefore analysed the performance of AFL when fewer models are generated to ensure a fairer comparison between the two methods. We find that reducing the AFL sampling to 500, 200, 100 or 50 models reduces the gain over AF2 to 1.30, 1.11, 0.86, and 0.63Å, respectively, compared to the above-reported average improvement of 1.53Å for 1000 models (Fig.3).

We also run AF2 for the antibodies in the LOW-80 dataset adding the antigen chain to the input data. This makes the AF2 calculations much slower, but we hypothesized that the presence of the antigen could improve the quality of the antibody loops, especially if the correct epitope has been identified. Across the LOW-80 dataset, we find that this is not the case, with the antigen-aware antibodies exhibiting an average H3-loop RMSD of 3.46±2.05Å, only slightly better than the antibody-only predictions, which feature an average H3-loop RMSD of 3.71±2.11Å. For this reason, we only consider antibody-only models in the remaining of the manuscript.

### 3.3 Clustered AlphaFlow ensembles

In the previous section, we have seen how AFL can generate predictions that outperform AF2 in terms of H3 loop accuracy for some difficult cases. Even reducing the number of generated models to a few tens allows us to retain a substantial improvement when considering the most accurate model in the AFL ensemble. Identifying such a model in a large pool of predictions is however extremely difficult. Here we show how structure-based clustering can help remove redundancy in the AFL ensemble, limiting the ensemble size at limited loss in prediction accuracy.

We took the AlphaFlow ensembles calculated over the LOW-80-W dataset and applied an Agglomerative Clustering procedure with three different linkage strategies (see Methods) and different numbers of clusters, K, ranging from 5 to 40. We observe that hierarchical clustering with complete and average linkage performs much better than single linkage when considering the quality of the best cluster center. When N=100 and K=20, complete linkage hierarchical clustering generates ensembles that are, on average, 1.38Å more accurate than the AF2 counterpart, compared to 1.27 and 0.72Å, respectively, for average and single linkage clustering.

Generating only 100 AFL models and clustering them with K=20 are reasonable choices to limit the computational time without losing too much H3 loop quality: with those settings, an AFL run takes approximately the same time as a standard AF2 run with five seeds (thus generating 25 models), allowing us to retain better conformations in the clustered ensemble, while still keeping the number of models relatively low, which can be beneficial in downstream tasks like docking.

### 3.4 Clustered AlphaFlow ensembles perform well Information-driven antibody-antigen docking

Having demonstrated a consistent improvement in H3 loop conformation RMSD with a clustered subset of the AFL ensemble, we now show that using these clustered ensembles as a starting point for docking outperforms the same protocol starting with the 5 clustered models obtained from AF2. To this end, we first generate full antibody ensembles by merging our heavy chain predictions with the top-ranked AF2 light chain model (see Methods), filtering out models showing backbone clashes between the two chains.

We then perform information-driven docking with HADDOCK3 using two information scenarios described in our previous publication (Giulini *et al*., 2024): the ideal “Para-Epi” scenario in which real knowledge of the paratope and epitope is used to define ambiguous interaction restraints (AIRs) to drive the modelling process, and the more realistic “CDR-VagueEpi” scenario in which surface-exposed CDR loops residues and a wider epitope definition on the antigen are used (see Methods). We dock two sets of antibody ensembles (AFL, AF2) to both experimentally resolved antigen structures (taken from the reference complexes, unbound-bound docking) and AF2-predicted antigen structures (unbound-unbound docking). The latter represents a more realistic scenario for docking while the former allows one to concentrate on the impact of the H3 loop conformation accuracy of the docking results.

Results for the Para-Epi docking scenario (see Fig. 4, top row) show how HADDOCK models obtained starting from AFL ensembles tend to be more accurate than those generated from AF2 ensembles. More specifically, the AFL protocol produces a good model ranked in position 1 for 67% (6 out of 9) and 78% (7 out of 9) of the cases when using the bound or unbound antigen, respectively. When considering the first 10 models, there is always a model of at least of acceptable quality. In contrast, runs starting from AF2 structural ensembles only provide acceptable predictions for 55% (5 out of 9) of the cases when targeting the bound antigen and 44% (4 out of 9) when using the unbound antigen. Importantly, even when these acceptable predictions exist, they are very rarely present within the first 10 models, thus making their identification challenging. AFL ensembles often yield medium-quality models, especially in the bound antigen scenario.

**Figure 4.**
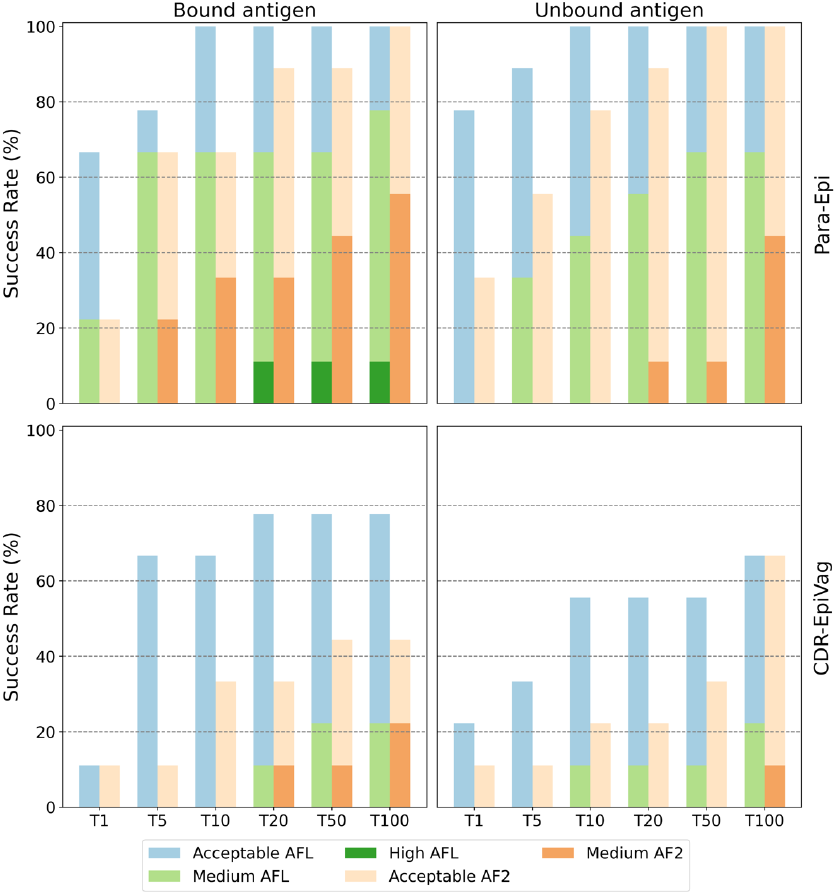
Success rate of HADDOCK runs using the Para-Epi (top row) and CDR-VagueEpi (bottom row) restraints to drive the docking. **Top row**: Runs starting from an AFL ensemble (blue/green colored-bars) always show at least an acceptable model within the top10, often yielding medium-quality models. In contrast, runs starting from AF2 ensembles (orange tints-colored bars) struggle to obtain good predictions, especially in the top 5-10 ranked models. **Bottom row:** AFL outperforms the AF2 protocol especially in the more realistic unbound-unbound docking scenario (bottom-right).

Our clustered diffusion approach also significantly outperforms AF2-based docking in the more realistic CDR-VagueEpi scenario, doubling the success rate on average. Considering the top 10 ranked models, our method achieves success rates of 67% for bound-unbound and 56% for unbound-unbound docking. The quality of the models is however lower, with only few medium quality predictions.

As a comparison, we also benchmarked the performance of AlphaFold3 (AF3) (Abramson *et al*., 2024) on the same dataset. The AF3 architecture contains a diffusion model that improves the diversity of the output models. Using the AF3 server, we generate a total of 25 models for each complex across 5 runs, each with a different random seed. Over the 9 cases where AF2 is not accurate (LOW-80-W dataset), AF3 outperforms AF2 in only 1 case, with RMSD of 2.1Å (7phu). Notably, the entire LOW-80-W dataset is included in the AF3 training set based on its training data cutoff. Overall, AF3 shows only slight improvements in H3 loop prediction compared to AF2. In terms of antibody-antigen complex prediction, similar to AF2-multimer, AF3 still struggles to produce acceptable or better models in the top10. For both tasks, increasing the number of used seed in AF3 is likely to provide more accurate predictions.

## 4 Conclusions

In this manuscript, we have presented an approach to improve the diversity of predicted antibody conformations and in particular of the H3 loop. We have demonstrated that AlphaFold2, in general, produces accurate antibody models, with the exception of the challenging H3 loop, which can be completely mispredicted. The H3 loop pLDDT serves as a good proxy for the accuracy of the prediction, allowing us to pinpoint the AlphaFold2 structures that resemble their experimental counterpart, but also identify cases where additional sampling and/or alternative approaches might be needed. In those cases, AlphaFlow provides highly heterogeneous ensembles that explore loop conformations that cannot be sampled through the standard AlphaFold2-multimer architecture, particularly when co-evolutionary signals are sparse. Unlike methods that rely heavily on MSA manipulation (Wayment-Steele *et al*., 2024; Stein and Mchaourab, 2022), AlphaFlow leverages flow matching to produce structurally diverse outputs without such dependencies. Structural ensembles from AlphaFlow can then be reduced in what we call clustered diffusion ensembles, namely a set of highly diverse structures that can be used for downstream tasks, such as docking. We have shown how the usage of these clustered diffusion ensembles dramatically improves the docking performance. A similar approach should also benefit the modelling of nanobodies, whose H3 loop is often even longer than in antibodies and, as such, even more challenging to model (Sircar *et al*., 2011; Gordon *et al*., 2023).

## Author contributions

Marco Giulini, Xiaotong Xu and Alexandre M.J.J. Bonvin designed the research. Marco Giulini and Xiaotong Xu conducted the research and analysed the results. All authors wrote and reviewed the manuscript.

## Supplementary data

Supplementary data are available online.

## Data availability

The input datasets and codes involved in this research are available at https://github.com/haddocking/alphaflow-antibodies. All the resulting modelling data are available from Zenodo (https://zenodo.org/records/14906314). The source code of HADDOCK3 is freely available at https://github.com/haddocking/haddock3.

## References

Abanades, B., Wong, W.K., Boyles, F., Georges, G., Bujotzek, A., and Deane, C.M. (2023) ImmuneBuilder: Deep-Learning models for predicting the structures of immune proteins. Commun Biol, 6, 575.

Abramson, J., Adler, J., Dunger, J., Evans, R., Green, T., Pritzel, A., Ronneberger, O., Willmore, L., Ballard, A.J., Bambrick, J., Bodenstein, S.W., Evans, D.A., Hung, C.-C., O’Neill, M., Reiman, D., Tunyasuvunakool, K., Wu, Z., Žemgulytė, A., Arvaniti, E., Beattie, C., Bertolli, O., Bridgland, A., Cherepanov, A., Congreve, M., Cowen-Rivers, A.I., Cowie, A., Figurnov, M., Fuchs, F.B., Gladman, H., Jain, R., Khan, Y.A., Low, C.M.R., Perlin, K., Potapenko, A., Savy, P., Singh, S., Stecula, A., Thillaisundaram, A., Tong, C., Yakneen, S., Zhong, E.D., Zielinski, M., Žídek, A., Bapst, V., Kohli, P., Jaderberg, M., Hassabis, D., and Jumper, J.M. (2024) Accurate structure prediction of biomolecular interactions with AlphaFold 3. Nature, 630, 493–500.

Baek, M., DiMaio, F., Anishchenko, I., Dauparas, J., Ovchinnikov, S., Lee, G.R., Wang, J., Cong, Q., Kinch, L.N., Schaeffer, R.D., Millán, C., Park, H., Adams, C., Glassman, C.R., DeGiovanni, A., Pereira, J.H., Rodrigues, A.V., Van Dijk, A.A., Ebrecht, A.C., Opperman, D.J., Sagmeister, T., Buhlheller, C., Pavkov-Keller, T., Rathinaswamy, M.K., Dalwadi, U., Yip, C.K., Burke, J.E., Garcia, K.C., Grishin, N.V., Adams, P.D., Read, R.J., and Baker, D. (2021) Accurate prediction of protein structures and interactions using a three-track neural network. Science, 373, 871–876.

Berman, H.M., Westbrook, J., Feng, Z., Gilliland, G., Bhat, T.N., Weissig, H., Shindyalov, I.N., and Bourne, P.E. (2000) The Protein Data Bank. Nucleic Acids Research, 28, 235–242.

Dominguez, C., Boelens, R., and Bonvin, A.M.J.J. (2003) HADDOCK: A Protein−Protein Docking Approach Based on Biochemical or Biophysical Information. J. Am. Chem. Soc., 125, 1731–1737.

Dunbar, J., Krawczyk, K., Leem, J., Baker, T., Fuchs, A., Georges, G., Shi, J., and Deane, C.M. (2014) SAbDab: the structural antibody database. Nucl. Acids Res., 42, D1140–D1146.

Giulini, M., Schneider, C., Cutting, D., Desai, N., Deane, C.M., and Bonvin, A.M.J.J. (2024) Towards the accurate modelling of antibody−antigen complexes from sequence using machine learning and information-driven docking. Bioinformatics, 40, btae583.

Gordon, G.L., Capel, H.L., Guloglu, B., Richardson, E., Stafford, R.L., and Deane, C.M. (2023) A comparison of the binding sites of antibodies and single-domain antibodies. Front. Immunol., 14.

Honorato, R.V., Trellet, M.E., Jiménez-García, B., Schaarschmidt, J.J., Giulini, M., Reys, V., Koukos, P.I., Rodrigues, J.P.G.L.M., Karaca, E., van Zundert, G.C.P., Roel-Touris, J., van Noort, C.W., Jandová, Z., Melquiond, A.S.J., and Bonvin, A.M.J.J. (2024) The HADDOCK2.4 web server for integrative modeling of biomolecular complexes. Nat Protoc, 19, 3219–3241.

Jing, B., Berger, B., and Jaakkola, T. (2024) AlphaFold Meets Flow Matching for Generating Protein Ensembles.

Jumper, J., Evans, R., Pritzel, A., Green, T., Figurnov, M., Ronneberger, O., Tunyasuvunakool, K., Bates, R., Žídek, A., Potapenko, A., Bridgland, A., Meyer, C., Kohl, S.A.A., Ballard, A.J., Cowie, A., Romera-Paredes, B., Nikolov, S., Jain, R., Adler, J., Back, T., Petersen, S., Reiman, D., Clancy, E., Zielinski, M., Steinegger, M., Pacholska, M., Berghammer, T., Bodenstein, S., Silver, D., Vinyals, O., Senior, A.W., Kavukcuoglu, K., Kohli, P., and Hassabis, D. (2021) Highly accurate protein structure prediction with AlphaFold. Nature, 596, 583–589.

Kenlay, H., Dreyer, F.A., Cutting, D., Nissley, D., and Deane, C.M. (2024) ABodyBuilder3: improved and scalable antibody structure predictions. Bioinformatics, 40, btae576.

McCoy, K.M., Ackerman, M.E., and Grigoryan, G. (2024) A comparison of antibody-antigen complex sequence-to-structure prediction methods and their systematic biases. Protein Sci, 33, e5127.

North, B., Lehmann, A., and Dunbrack, R.L. (2011) A New Clustering of Antibody CDR Loop Conformations. Journal of Molecular Biology, 406, 228–256.

Raouraoua, N., Mirabello, C., Véry, T., Blanchet, C., Wallner, B., Lensink, M.F., and Brysbaert, G. (2024) MassiveFold: unveiling AlphaFold’s hidden potential with optimized and parallelized massive sampling. Nat Comput Sci, 4, 824–828.

Regep, C., Georges, G., Shi, J., Popovic, B., and Deane, C.M. (2017) The H3 loop of antibodies shows unique structural characteristics. Proteins, 85, 1311–1318.

Ruffolo, J.A., Chu, L.-S., Mahajan, S.P., and Gray, J.J. (2023) Fast, accurate antibody structure prediction from deep learning on massive set of natural antibodies. Nat Commun, 14, 2389.

Sircar, A., Sanni, K.A., Shi, J., and Gray, J.J. (2011) Analysis and Modeling of the Variable Region of Camelid Single-Domain Antibodies. The Journal of Immunology, 186, 6357–6367.

Stein, R.A. and Mchaourab, H.S. (2022) SPEACH_AF: Sampling protein ensembles and conformational heterogeneity with Alphafold2. PLOS Computational Biology, 18, e1010483.

Wallner, B. (2023) AFsample: improving multimer prediction with AlphaFold using massive sampling. Bioinformatics, 39, btad573.

Wayment-Steele, H.K., Ojoawo, A., Otten, R., Apitz, J.M., Pitsawong, W., Hömberger, M., Ovchinnikov, S., Colwell, L., and Kern, D. (2024) Predicting multiple conformations via sequence clustering and AlphaFold2. Nature, 625, 832–839.

Weitzner, B.D., Dunbrack, R.L., and Gray, J.J. (2015) The Origin of CDR H3 Structural Diversity. Structure, 23, 302–311.

Wu, J., Wu, F., Jiang, B., Liu, W., and Zhao, P. (2022) tFold-Ab: Fast and Accurate Antibody Structure Prediction without Sequence Homologs. 2022.11.10.515918.

