## supplementary material for "Improved structural modelling of antibodies and their complexes with clustered diffusion ensembles"

M. Giulini[1†], X. Xu[1†], A.M.J.J. Bonvin [1]*

[1]Computational Structural Biology Group, Department of Chemistry, Bijvoet Centre, Faculty of Science, Utrecht University

† These authors contributed equally to this work.

### Supplementary Table 1. CDR loop sequences of the 54 antibodies present in the data set

| PDB | H1 | H2 | H3 | L1 | L2 | L3 |
|---|---|---|---|---|---|---|
| 7bbj | GYTFTTYW | IYPGLSDT | ARLLDYAMDY | QDIRSY | YTS | QQGETLPWT |
| 7bnv | GYSFTSYW | IYPGDSDT | ARHPSPYYYGSGSYSGGFDY | QSVSSSY | GAS | QQYDNWPLMHT |
| 7daa | GFSLSDYA | IYASGST | ARYYAGSDI | QSISAY | DAS | QTYYAIITYGAA |
| 7dk2 | GFTFSSYW | IKQDASEK | ARDLGILWFGDYP | QGISNS | AAS | QQFYSTPRT |
| 7e72 | GYSFTSYW | IHPSDSET | ARGLYGNS | QDIGIS | ATS | LQYASSPYT |
| 7f7e | GFTFSSYA | IVGSGGST | AKSLIYGHYDILTGAYYFDY | QGIGNW | AAS | QQANSFPP |
| 7kez | GFNIKDTY | IYPTNGYT | ARGGAVAGTGVYYFDY | QDIPRSISGY | WGS | QQHYTTPPT |
| 7kf0 | GFNIKDTY | IYPTNGYT | ARGGSFYYYYMDV | QDIPRSISGY | WGS | QQHYTTPPT |
| 7kf1 | GFNIKDTY | IYPTNGYT | AKLGIGYYYYGMDV | QDIPRSISGY | WGS | QQHYTTPPT |
| 7kql | GGSISSRSYY | IYYSGFT | ATGGPYGDYAHWFEP | QSVSSSY | GAS | QQYGSSPIT |
| 7l7r | GFTFSSYV | IYRGGST | VKDPKAWLEPEW | QSISKY | AAS | QQSYSNPRT |
| 7lr3 | EFTFSDYG | ISSGSNSI | SREAYFAMDY | QDIHKY | YTS | LQYDNLYT |
| 7lr4 | DYSLSDYN | INPNHGTT | ASPIHYGNHVPFDY | QDISNY | FTS | QQGITLPWT |
| 7mdj | GYTFTSDW | IIPSYGRA | ARERGDGYFDY | QSIGTD | YAS | QQSNRWPFT |
| 7mrz | GGSISSSSYY | ISYSGST | ARDSLRYGMDV | QSVLYSSNNKNY | WAS | QQYALAPPRT |
| 7msq | GGSISSYH | IYYSGNT | VREMRRGYSGYDYWDLYAFDI | QGISSY | AAS | QQLNSYPHT |
| 7mzf | EFIVSRNY | IYSGGTT | ARDRGDYLFDY | QSISSW | KAS | QQYNSYFPT |
| 7mzg | GLTVSSNY | FYPGGST | ARDAVYYGMDV | QSISSY | AAS | QESYSTPGLFT |
| 7mzh | GYTFTGYY | INPNSGGT | ARSYYDY | SSNIGHNA | YDD | AAWDDILNGPV |
| 7mzi | GFTFSRFA | ISGSGGST | AKVGWGAFDI | YSNIGSNP | AND | STWDDSLPGPL |
| 7mzj | GFTFSYAW | IKRKSDGGTT | TTDLCRSTSCEHDAFDI | QSIRSY | AAS | QQSYTTPAIT |
| 7mzk | GYTFTSYY | INPSGGGT | AKDRVTIFWGNGMDV | QSVLYSSNNKNY | WAS | HQYSTTPLT |
| 7n4i | GYSFISYW | IYPGDSDT | ARLLYYSDSSPLDS | QSISTY | AAS | QQSHSTPRT |
| 7n4j | GDSISSSDYS | IYYIKNT | ARERPPFDVVVVPAARPYNWFDP | SSNIGAGYD | GNN | QSYDSSLSGSKV |
| 7np1 | GITVSSNY | IYSGGST | ARGEGGSIVGVTSDY | QSISRY | AAS | QQSYSTLPYT |
| 7nx3 | GYAFSSYW | IYPGDGDT | ARSRGYFYGSTYDS | ESVDNYGISF | AAS | QQSKEVPWT |
| 7phu | GYTFTDYY | INPNSGGT | ARDLWFGESPPYYGVDV | NIGSYS | YDS | QVWDTNTDHVV |
| 7phw | GFDFSSYA | IRNDGSFT | TKSADDGGHYSDFSGEIDA | TYNY | YND | GNSDSRNVA |
| 7pi7 | GFNIKDTY | IDPANGNT | ARDVLYFDV | ESVDSYGNSF | RAS | QQSNEDRT |
| 7pqy | ASGFTVSSNYMS | SVIYSGGSTYY | YCARDHVRPGMNIWG | CQASQDISNY | LLIYDAS | ATYYCQQYDNLPV |
| 7pr0 | GFTFSSYS | ISSSSSTI | ASPGGITAAGTSVLFGYYGMDV | QSLLHSNGYNY | LGS | MQALQTPITWT |
| 7ps0 | DGSISSSDYY | IYYTGST | ARLVVPSPKGSWFDP | SIDVGNYNL | EGS | CSYVGSSTYV |
| 7ps1 | GLTVRSNY | IYSGGST | ARDLVVYGMDV | QSVSSS | GTS | QQYGSSPL |
| 7ps2 | GFTFSNYG | ISYEESNR | AKDQGPATVMVTAIRGAMDV | QSVLYSSNNKNY | WAS | QQYFGSPSIT |
| 7ps4 | GYSFTNYW | IYPGDSGT | ARSRVGATGGYYDYYMDV | SSNLGGNT | SNN | AAWDDSLNGPV |
| 7ps6 | GGSITSSNHY | MYYSGST | ARQIGPKRPSQVADWFDP | QGISSY | AAS | QQLNSYPLT |
| 7q0g | GGTFSSSV | IIPLFGSA | AKVSQWALILF | QSVSSSY | GAS | QQYGTSPSWT |
| 7q0i | GFTFSSYG | IWYDGSNN | ARSYCSGGFCFGYYYGLDV | NIGTKS | YNS | QVWDSGSDHYV |
| 7qnw | GDSISSSRYY | FYYSGIT | ARPRPPDYYDNSGALLFDI | QSISAW | KAS | QQYISSSPWT |
| 7qny | GFTFDDYA | VSWNSGTI | AREVGGTFGVLISREGGLDY | TIGSKS | DDS | QVWDSSSDRVV |
| 7qu1 | GYAFGSHW | IYPGDGDT | ARDDYGTRYYFDY | QDINNY | YTS | QQGKTLPLT |
| 7qu2 | GFTFSNYQ | ITVKSDNYGA | SRSGIYDGYYAYAMDY | QIVGTS | WAS | QQYATYPLT |
| 7rfb | GGTVNT | IFPLLGVP | AKDGVGWSGHGPSQWSGVDV | QSLLHSTGYNY | LGS | MQALEIPRLT |
| 7s0b | GFTFSSYA | ISGSGGST | ARDLWGSGFFAFDV | QDISNY | DAS | QQDAGTPLT |
| 7s11 | GLSLTTNS | IWSNGGT | ARNFPYPGINFD | TGAVTTSNY | GTS | SLWYSGHLI |
| 7s13 | GLSLTNNI | IWSNGGT | ASRDYPGFAY | ELPKRY | EDS | LSTYSDDKLPI |
| 7seg | GYTFTSYY | IEPMYGST | ARGSAYYYDFADY | NIGSKN | QDN | QVWDNYSVL |
| 7sem | GFTFSSYS | ISASSSYS | ARARATGYSSITPYFDI | SSNIGAGYD | DNN | QSYDRSLSGV |
| 7shu | GYNITSGYS | VTYDGST | AKGNNYFGHWHFAV | KSVDSDGDSY | AAS | QQSHEDPYT |
| 7shz | GYSITSGYS | IKYSGET | ARGSHYFGHWHFAV | KPVDGEGDSY | AAS | QQSHEDPYT |
| 7si0 | GYSITSGYS | VTYDGST | ARGSHYFGHWHFAV | QSVDSDGDSY | AAS | QQSHEDPYT |
| 7so9 | ASGFTFNSYGMH | AFIRYDGGNKYY | YCANLKDSRYSGSYYDYWG | CQASQDIRFY | LLISDAS | ATYYCQQYDNLPF |
| 7stz | VSGFSLSRYGVH | GMMWGGGNTDY | YCASSNYVLGYAMDYWG | CKSSQSLLNSSNQKNY | LLIYFTS | ADYFCQQHYRTPH |
| 7vux | GFAFSSYD | ISGGGRYT | ASPYGGYFDV | QSISNF | YAS | QQSNSWPHT |

### Supplementary Figure 1. Best ranked loops in terms of loop PLDDT

We here report the RMSD values of the loop with the highest H3 loop pLDDT. Although the latter is a local measure tailored to predict the quality of the loop, it seems to perform slightly worse than the global Alphafold2 ranking score.

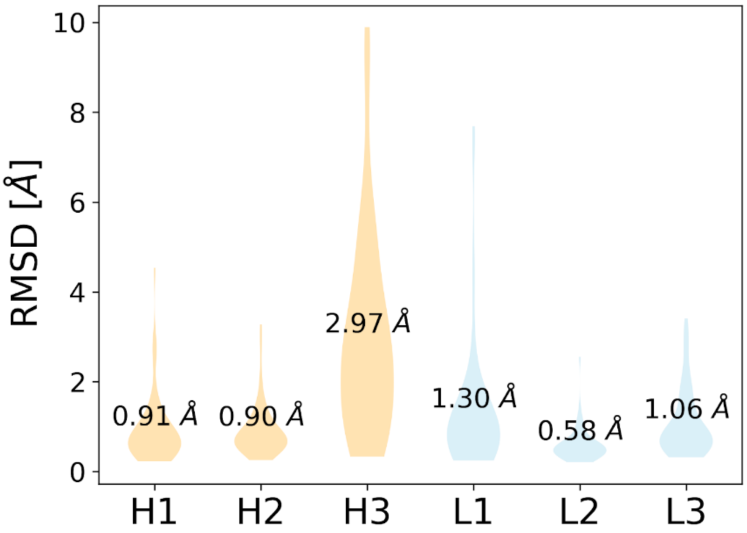


**SI Figure 1.** Violin plot of loop accuracy (measured by the loop RMSD from the reference crystal structure after superimposition on the frame-work region) over the six antibody hypervariable loops for the loops with the highest H3 loop pLDDT.

#### Supplementary information 1. An example HADDOCK configuration file for docking

```
# ======================================================
# Example antibody-antigen HADDOCK docking workflow
# ======================================================

# run directory
run_dir = "antigen-AFL-Para-Epi-full-1000-200"
mode = "local"
ncores = 24
clean = true

#Input proteins
molecules =  [
    "ensemble_PDB_emref.pdb",
    "PDB_antigen_haddock-ready.pdb"
    ]
# ======================================================
# Parameters for each stage of the workflow
# ======================================================

[topoaa]

[rigidbody]
# CDR to surface ambig restraints
ambig_fname = "PDB_ambig_Para_Epi.tbl"
# Restraints to keep the antibody chains together
unambig_fname = "PDB_unambig_AF2.tbl"
sampling = 1000

[caprieval]
reference_fname = " PDB_target.pdb"

[seletop]
select = 200

[flexref]
tolerance = 20
# CDR to surface ambig restraints
ambig_fname = "PDB_ambig_Para_Epi.tbl"
# Restraints to keep the antibody chains together
unambig_fname = "PDB_unambig_AF2.tbl"

[emref]
tolerance = 5
# CDR to surface ambig restraints
ambig_fname = PDB_ambig_Para_Epi.tbl"
# Restraints to keep the antibody chains together
unambig_fname = “PDB_unambig_AF2.tbl"

[caprieval]
reference_fname = “PDB_target.pdb"

[clustfcc]
min_population=4

[caprieval]
reference_fname = “PDB_target.pdb"

# ======================================================
```

**Note** that in a real case, the reference structure won't be known and the "`reference_fname`" lines in the "`caprieval`" modules should be removed or commented out.
